# PHESANT: a tool for performing automated phenome scans in UK Biobank

**DOI:** 10.1101/111500

**Authors:** Louise A C Millard, Neil M Davies, Tom R Gaunt, George Davey Smith, Kate Tilling

## Abstract

**Motivation:** Epidemiological cohorts typically contain a diverse set of phenotypes such that automation of phenome scans is non-trivial, because they require highly heterogeneous models. For this reason, phenome scans have to date tended to use a smaller homogeneous set of phenotypes that can be analysed in a consistent fashion. We present PHESANT (PHEnome Scan ANalysis Tool), a software package for performing comprehensive phenome scans in UK Biobank.

**General features:** PHESANT tests the association of a specified trait with all continuous, integer and categorical variables in UK Biobank, or a specified subset. PHESANT uses a novel rule-based algorithm to determine how to appropriately test each trait, then performs the analyses and produces plots and summary tables.

**Implementation:** The PHESANT phenome scan is implemented in R. PHESANT includes a novel Javascript D3.js visualization, and accompanying Java code that converts the phenome scan results to the required JavaScript Object Notation (JSON) format.

**AVAILABILITY:** PHESANT is available on GitHub at [https://github.com/MRCIEU/PHESANT]. Git tag v0.2 corresponds to the version presented here.

## INTRODUCTION

Phenome scans test the association of a trait of interest with a comprehensive array of phenotypes (the “phenome”). Types of phenome scans include phenome-wide association studies (pheWAS) (1), Mendelian randomization-pheWAS (MR-pheWAS) (2) and environment-wide association studies (EnWAS) (3,4). PheWAS seek to investigate the association of a genetic variant with a set of phenotypic traits (1,5). A recent extension to pheWAS, MR-pheWAS, uses Mendelian randomization (MR) in a pheWAS framework in order to search for the causal effects of a particular exposure (2). EnWAS seek to test the associations of a trait of interest with a set of other phenotypes (3).

Epidemiological cohorts usually contain a large number of diverse phenotypes, such that testing the association of these phenotypes with another trait in an automated way is nontrivial. For this reason, researchers wishing to perform a phenome scan will typically specify a homogeneous subset of traits, in order to automate the tests of association across these traits in a consistent way. For instance, pheWAS initially started using international classification of disease (ICD) codes from electronic health records, where each disease code could be treated as a binary variable and a consistent test performed (5). However, restricting the set of phenotypes provides only a partial view of associations with a trait of interest, and reduces the potential to identify novel associations.

In this paper we present PHESANT (PHEnome Scan ANalysis Tool), a parallelizable tool for phenome scans in UK Biobank, a prospective cohort of over 500 000 men and women in the UK aged between 37–73 years (6). This cohort includes genetic data, and a large and diverse range of data from blood, urine and saliva samples analyses, clinical assessments, record linkage and health and lifestyle questionnaires. The diversity of traits available coupled with the large sample size provides an opportunity to identify novel associations with phenome scans.

## IMPLEMENTATION

PHESANT is implemented in R and requires the following R packages: optparse, MASS, lmtest, nnet and forestplot (see GitHub repository for package versions). PHESANT takes one data file as input containing the set of phenotypes and the trait of interest, which may be a SNP, a genetic score or a phenotypic trait depending on whether a pheWAS, MR-pheWAS or EnWAS is being performed (the trait of interest can alternatively be provided as a separate file if this is preferred). PHESANT also makes use of two data files that contain information about the variables in the UK Biobank cohort: 1) a data coding information file, and 2) a variable information file. These files have been set up for the example we describe in the usage section, but can be changed as needed for each particular phenome scan. For more information on the PHESANT data and information files see the documentation in the GitHub repository. In the rest of this section we describe the variable processing flow used in PHESANT.

### Automated processing flow to determine variable coding

In order to test the association of the trait of interest with the diverse range of phenotypes in UK Biobank in an automated manner, we developed a rule-based system to determine the appropriate coding of each variable and hence test of association to use. These rules are shown in Figure 1 and described in full in the Supplementary section S1. The decision rules start with the variable *field* type (as specified by UK Biobank at [http://biobank.ctsu.ox.ac.uk/showcase/list.cgi]), either continuous, integer, categorical (single) or categorical (multiple), and categorise each variable as one of four *data* types: continuous, ordered categorical, unordered categorical and binary. The categorical *(single)* field type refers to categorical fields (including binary) where each participant can only have one value. For example, by questionnaire participants were asked “How would you describe your usual walking pace?” with options including “slow”, “average” and “brisk” (field ID=924; see Supplementary figure 1). In contrast, categorical *(multiple)* fields can have multiple values per participant. For example, by questionnaire participants were asked what types of bread they ate the previous day (field ID=20091; see Supplementary figure 2) and could, for instance, select both white and wholemeal options. Where a field is measured at several time points we use the first occurrence only (see Supplementary section S2 for details). Continuous and integer variables may have more than one measurement at this first measured time point (typically to improve the estimate of a measurement). For instance, spirometry was measured two or three times a few moments apart (see for example field 3062 [http://biobank.ctsu.ox.ac.uk/showcase/field.cgi?id=3062]). When this is the case we take the mean to create a single value per participant (see Supplementary section S2 for details).

**Figure 1:**
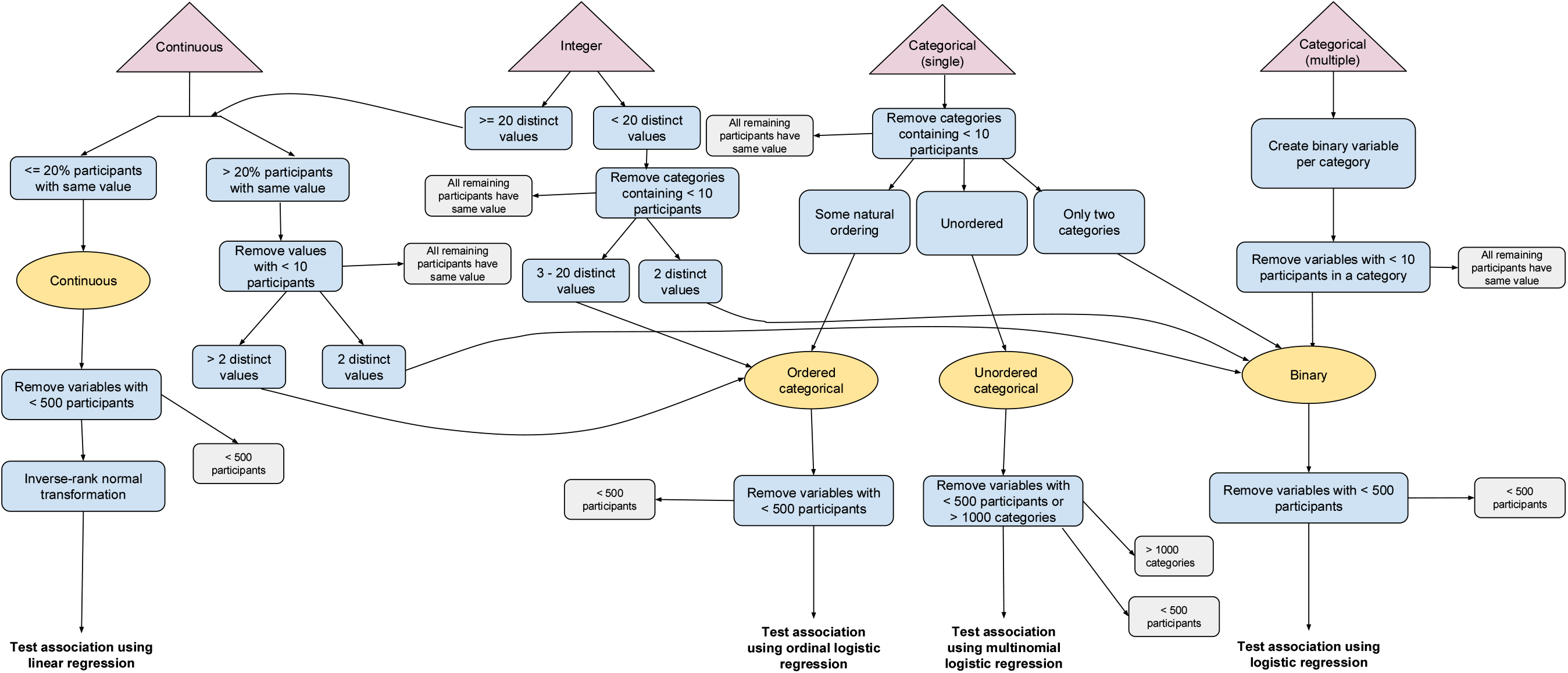
Variable processing flow diagram showing logic from defined field type specified by UK Biobank to test of association. Pink triangular nodes at top of figure are field types defined by UK Biobank. Blue rectangular nodes show processing logic used to determine the data type assignment (yellow, oval) – either: continuous, ordered categorical, unordered categorical and binary, and hence finally, the type of test used: linear, ordinal logistic, multinomial logistic and logistic regression, respectively. Grey rectangular nodes show points where variables may be removed.

Variables with the continuous field type are usually assigned to the continuous data type. In this case, the variable is transformed to a normal distribution using an inverse normal rank transformation. In a minority of cases, continuous fields are assigned to the ordered categorical data type (or binary if there are only two distinct values). For example, field 100022 [http://biobank.ctsu.ox.ac.uk/showcase/field.cgi?id=100022] contains the estimated alcohol intake based on responses to the “diet by 24-hour recall” questionnaire. A large proportion of the participants have a zero value for this field, because they consumed no alcohol. It is not possible to inverse normal rank transform this variable because where a large number of participants have the same value the rank assigned in this transformation is random among these, and this would add noise to the data. Instead, we transform this variable into three categories with roughly the same number of participants in each (placing split points between distinct values), and treat it as an ordered categorical variable (see algorithm in Supplementary section S3). Variables with the integer field type are usually treated exactly the same as the continuous variables. In a minority of cases, where there are 20 or fewer distinct values we treat this variable as ordered categorical (or binary if there are only two distinct values).

Categorical (single) variables may be assigned to the binary, ordered categorical or unordered categorical data types. UK Biobank consistently assigns negative values to categories denoting missingness (such as “Preferred not to answer” and “Do not know”), and so we recode negative values to NA. UK Biobank defines ‘data codes’ to which one or more fields are assigned, and these define the set of categorical values for these fields and their corresponding numeric values. The PHESANT data-coding information file specifies whether a data code of a categorical (single) field defines an ordered or unordered category structure, and we use this information to assign each non-binary categorical (single) field as either an ordered or unordered categorical data type.

Each categorical (multiple) variable is converted to a set of binary variables, each denoting whether a participant has a given value of this variable. For example, for the variable describing the bread eaten yesterday (field ID=20091; Supplementary figure 2), with values ‘white’, ‘mixed’, ‘wholemeal’, ‘seeded’ and ‘other’, we generate 5 binary variables, white={true,false}, wholemeal={true,false} and so forth. Categorical (multiple) fields have an added complexity because when a person has no value in this field this may be because: 1) the field values are *incomplete* – they do not contain all possible values (e.g. a participant who does not eat bread cannot choose any option above) or 2) because the data is missing (e.g. because a participant did not answer this particular question). This affects who we assign as, for instance, ‘white=false’, either 1) all people who selected a value other than ‘white’, 2) all people who responded to this questionnaire and did not select ‘white’ or 3) all people who did not select ‘white’ including those who did not respond to the questionnaire (see Supplementary figure 3 for illustration). In this case the second option might be preferred such that we are comparing those who ate white bread with those who responded to the questionnaire but did not eat white bread. This decision is variable specific, and can be specified in the PHESANT variable information file (the variable information file we used for the example described in the next section is available in the GitHub repository). For more details see Supplementary material section S1.

Some categorical multiple fields include negative numeric values for particular categories denoting missingness (such as “Do not know”). We exclude all participants with a missing value from the *false* value of the generated binary variable, because we cannot know if they do or do not pertain to the *true* value of this binary variable. For example, consider field 41228 [http://biobank.ctsu.ox.ac.uk/showcase/field.cgi?id=41228] describing the type of medical professional who conducted the delivery of a participant’s child, and a participant who has given birth twice and has values “midwife” and “not known” in this field. The generated binary variable for midwife includes this participant in the set of participants corresponding to *midwife=true* because we know that on at least one occasion a midwife conducted the delivery. However, we cannot be certain that a hospital doctor did not conduct a delivery for this participant because the “not known” value could refer to a “hospital doctor”. Hence, the generated “hospital doctor” binary variable would not include this participant in the set of participants corresponding to *hospital_doctor=false,* because this is not known.

### Tests of association with trait of interest

The association of each phenotype, having been appropriately coded and assigned one of the four data types (continuous, ordered categorical, unordered categorical and binary), is tested with the trait of interest as follows. The phenotype and the trait of interest are the dependent and independent variables of the regression, respectively. All regressions are adjusted for age at recruitment and sex (and also genotype chip when the trait of interest is genetic, derived from the genotype measurement batch). We test the association with the transformed variables of the continuous data type using linear regression (lm R function). Ordered categorical, unordered categorical and binary variables are tested using ordered logistic regression (polr R function), multinomial logistic regression (multinom R function) and binomial regression (glm R function with family parameter as binomial), respectively. We do not test phenotypes where the sample size is fewer than 500, which typically occurs for a minority of fields such as follow up questions on a subsample (e.g. field 22148 [http://biobank.ctsu.ox.ac.uk/showcase/field.cgi?id=22148]). We do not test unordered categorical variables with more than 1000 categories (above the default maximum of the multinom function), which occurs once in our usage example (for field 132 [http://biobank.ctsu.ox.ac.uk/showcase/field.cgi?id=132]).

### Customising a phenome scan with PHESANT

PHESANT allows researchers to easily customise the phenome scan by changing settings in the data coding and variable information files. This includes:

- Changing the numeric values underlying a variable (such as recoding a value to missing) or the ordering of values for ordered categorical variables.
- Assigning a default value to categorical (single) variables where this is not explicitly coded in the variable (see Supplementary section S1).
- Changing fields from the categorical (single) to the categorical (multiple) field type, as this may be more appropriate for a small number of fields.
- Specifying which variables should be excluded a priori from the phenome scan.
- Specifying which fields in the phenome dataset are essentially the same phenotype as the trait of interest (e.g. weight and body mass index (BMI)), such that, after the phenome scan is run, the results of association between these fields and the trait of interest are used for validation only, rather than being included in the results and adding to the multiple testing burden.

### PHESANT-viz: a web-based visualization for phenome scans

Reviewing the results from phenome scans can be challenging due to the number and complexity of phenotypes. As part of PHESANT we have also developed PHESANT-viz, a D3 Javascript visualization that displays phenome scan results as an interactive graph, using the hierarchical field category structure defined by UK Biobank (available at [http://biobank.ctsu.ox.ac.uk/showcase/label.cgi]). PHESANT includes a Java program to convert the phenome scan results to the JavaScript Object Notation (JSON) format required for PHESANT-viz. We provide the PHESANT-viz of our usage example below.

## USAGE

To demonstrate PHESANT we have performed a MR-pheWAS, to search for the causal effects of BMI in UK Biobank (previously performed in a smaller cohort with continuous phenotypes only (2)). Such an analysis is predicated on the Mendelian randomization principle that genetic variants can be used as instrumental variables to estimate causal effects of the phenotype they proxy for on downstream outcomes (7). In the current context this would be a screening exercise to identify associations for detailed follow-up. This analysis is preliminary and for example only, having been run on a non-random subsample of 114 963 participants (containing the UK BILEVE samples selected on smoking status; see Supplementary section S4 for details) for which genetic data is currently available in UK Biobank (a final analysis will be subsequently published upon release of the full 500 000 sample with genetic data).

We created an allele score from 96 genetic variants previously found to be associated with BMI, in a recent genome-wide association study (GWAS) meta-analysis (8). The score was calculated as a sum of the number of BMI-increasing alleles, weighted by the effect size as reported in (8) (see Supplementary table 1). Hence, a higher genetic score corresponds to a tendency towards higher BMI (F-statistic=1979). We used PHESANT to test the association of the BMI genetic score with the 290 integer, 1030 continuous, 658 categorical (single) and 99 categorical (multiple) fields available in UK Biobank at the current time (excluding 66 fields a priori, see Supplementary table 2). Supplementary figure 4 shows the number of variables reaching each stage of our variable processing flow.

Figure 2 shows the QQ plot of our MR-pheWAS results (see full results ranking in Supplementary data table, and forest plots in Supplementary figure 5). Of the 12 819 tests performed (excluding 87 phenotypes tested but specified a priori as being aspects of the same essential phenotype as BMI), 86 were associated at a Bonferroni corrected P value threshold of 3.90×10^-6^ (0.05/12819). We detected several known effects of BMI, for example with hypertension (9) (fields 41204 value I10, and 4079), diabetes (10) (field 2443) and age at puberty in both sexes (11) (fields 2714, 2375 and 2385). For instance, a 1 standard deviation (SD) increase in BMI allele score was associated with a 1.09-fold [95% confidence interval (CI): 1.06, 1.11] higher odds of being diagnosed with hypertension in hospital (field 41204 value I10), and a 0.015 SD [95% CI: 0.010, 0.021] higher diastolic blood pressure (field 4079). We also detected a number of potentially causal associations that were previously unknown. For example, participants with a genetic propensity to higher BMI were less likely to perceive themselves as a nervous person (field 1970) or to call themselves tense or ‘highly strung’ (field 1990).

The PHESANT-viz of these preliminary results can be found at [datamining.org.uk/PHESANT/] or within the PHESANT package. When this analysis is performed using the full 500 000 sample the power to detect associations will increase, and it is also likely that the number of tests will increase as fewer variables will be filtered out in the variable processing steps (due to a small sample size). This preliminary analysis took approximately 81 hours (using a 1 core Intel E5-2670).

**Figure 2:**
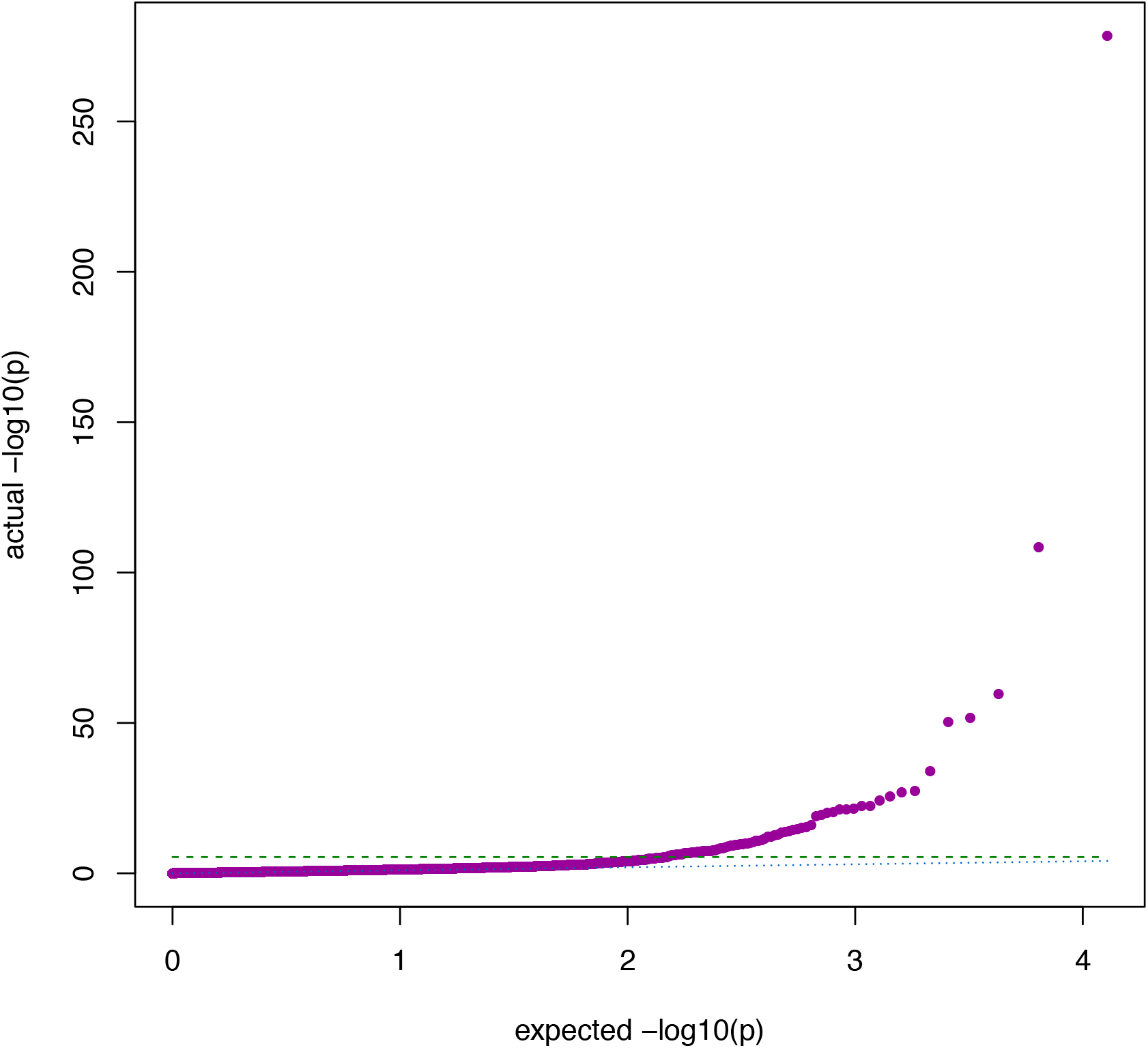
QQ plot of preliminary MR-pheWAS analysis seeking to identify the causal effects of BMI. Green dashed line: Bonferroni corrected threshold p= 3.90 x 10^-6^ (p=0.05 corrected for 12 819 tests). Blue dotted line: actual = expected.

## DISCUSSION

PHESANT enables researchers to perform a comprehensive phenome scan in UK Biobank, including a pheWAS, MR-pheWAS or EnWAS. While GWAS have been highly successful at identifying novel associations, we are currently only just beginning to explore the phenome in a hypothesis-free manner (1,2,12). In contrast to hypothesis-driven analyses, phenome scans allow exploration across hypotheses without strong priors, and should help to avoid publication bias as analyses are pre-specified and all results, not just the most ‘statistically significant’, are published together.

When undertaking a phenome scan there are several important considerations. First, phenome scans are a screening exercise to identify potentially interesting associations that should then be analysed more rigorously - and as such the effect estimates should be interpreted with caution. The strength of the strongest associations identified may be inflated due to the winners curse. Second, it is important to consider the number of tests performed when examining the strength of identified associations. In our usage example we used a conservative Bonferroni corrected threshold to identify potentially interesting associations, which, while reducing the type I error rate is likely to increase the type II error rate. Third, interpretation of potentially hundreds of results is challenging because the correlated structure of phenotypes means that associations between the trait of interest and each phenotype are not independent. The strongest associations should not be viewed in isolation but alongside the results of related variables for which an association may not have been identified, and to do this we provide a novel visualization approach, PHESANT-viz.

We note the following limitations and areas of future work. It is possible that in some cases our automated rule-based method may deal with variables inappropriately. For example, field 132 [http://biobank.ctsu.ox.ac.uk/showcase/field.cgi?id=132], which describes participant’s jobs, is treated as an unordered categorical variable but was removed from our phenome scan because it has more than 1000 categories. In this case, it may be preferable to use the hierarchical structure defined by UK Biobank to combine categories of related jobs.

PHESANT uses the first time point of a field where multiple time points are available. In future work we will investigate how to automate the analysis including data from multiple time points and the potential gains that this would give (13). Currently, when continuous variables are converted to the ordered categorical data type we arbitrarily chose to generate three categories, and in future work with will investigate whether a larger number of categories are beneficial, and whether the optimal number of categories can be calculated from the distribution of the variable. PHESANT is specifically designed for use in UK Biobank, but a cohort-independent tool for phenome scans would be highly valuable. Hence, in the future we will aim to adapt PHESANT for use in a general setting. Finally, we intend to integrate PHESANT with MR-base (14), to enable automated construction of genetic instrumental variables to use in MR-pheWAS.

The large number of participants combined with the extensive range of phenotypes available in UK Biobank provides a great opportunity to comprehensively search for novel (potentially causal) associations in a hypothesis-free manner. We are aware of only one very small phenome scan that has been performed in UK Biobank to date (15), and a recent novel Bayesian approach of self-reported diagnoses and hospital episodes (16). To our knowledge, PHESANT is the first open source package to automate phenome scans across diverse sets of phenotypes.

## REFERENCES

1. Denny J, Bastarache L, Roden DM. Phenome-Wide Association Studies as a Tool to Advance Precision Medicine. Annu Rev Genomics Hum Genet. 2016;17:353–73.

2. Millard LAC, Davies NM, Timpson NJ, Tilling K, Flach PA, Davey Smith G. MR-PheWAS: hypothesis prioritization among potential causal effects of body mass index on many outcomes, using Mendelian randomization. Sci Rep. Nature Publishing Group; 2015 Jan 16;5:16645.

3. Tzoulaki I, Patel C, Okamura T, Chan Q, Brown I, Miura K, et al. A nutrient-wide association study on blood pressure. Circulation. 2012;126(21):2456–64.

4. Patel CJ, Bhattacharya J, Butte AJ, Schwartz D, Collins F, Hirschhorn J, et al. An Environment-Wide Association Study (EWAS) on Type 2 Diabetes Mellitus. Zhang B, editor. PLoS One. Public Library of Science; 2010 May 20;5(5):e10746.

5. Denny J, Ritchie M, Basford M, Pulley J, Bastarache L, Brown-Gentry K, et al. PheWAS: demonstrating the feasibility of a phenome-wide scan to discover gene-disease associations. Bioinformatics. 2010;26(9):1205–10.

6. Allen N, Sudlow C, Downey P, Peakman T, Danesh J, Elliott P, et al. UK Biobank: Current status and what it means for epidemiology. Heal Policy Technol. 2012; 1(3): 123–6.

7. Davey Smith G, Ebrahim S. “Mendelian randomization”: can genetic epidemiology contribute to understanding environmental determinants of disease? Int J Epidemiol. 2003 Feb;32(1):1–22.

8. Locke A, Kahali B, Berndt S, Justice A, Pers TH, Day FR, et al. Genetic studies of body mass index yield new insights for obesity biology. Nature. 2015;518(7538):197–206.

9. Timpson N, Harbord R, Davey Smith G, Zacho J, Tybjærg-Hansen A, Nordestgaard BG. Does greater adiposity increase blood pressure and hypertension risk? Mendelian randomization using the FTO/MC4R genotype. Hypertension. 2009;54(1):84–90.

10. Holmes M, Lange L, Palmer T, Lanktree M, North KE, Almoguera B, et al. Causal effects of body mass index on cardiometabolic traits and events: a Mendelian randomization analysis. Am J Hum Genet. 2014;94(2):198–208.

11. Mumby H, Elks C, Li S, Sharp S, Khaw K-T, Luben RN, et al. Mendelian randomisation study of childhood BMI and early menarche. J Obes. 2011;2011.

12. VanderWeele T. Outcome-wide Epidemiology. Epidemiology. 2017;

13. Cao Y, Rajan SS, Wei P. Mendelian randomization analysis of a time-varying exposure for binary disease outcomes using functional data analysis methods. Genet Epidemiol. 2016 Dec;40(8):744–55.

14. Hemani G, Zheng J, Wade KH, Laurin C, Elsworth B, Burgess S, et al. MR-Base: a platform for systematic causal inference across the phenome using billions of genetic associations. bioRxiv. 2016;

15. Emdin C, Khera A, Natarajan P, Klarin D, Won H-H, Peloso GM, et al. Phenotypic characterization of genetically lowered human lipoprotein (a) levels. J Am Coll Cardiol. 2016;68(25):2761–72.

16. Cortes A, Dendrou C, Motyer A, Jostins L, Vukcevic D, Dilthey A, et al. Bayesian analysis of genetic association across tree-structured routine healthcare data in the UK Biobank. bioRxiv. 2017;

